# *Lactobacillus rhamnosus* (LR) ameliorates acute respiratory distress syndrome (ARDS) via modulating the lung “Innate Lymphoid Cells (ILCs)-Mononuclear Phagocytic System (MPS)”

**DOI:** 10.1101/2023.08.10.552899

**Authors:** Leena Sapra, Sneha Das, Chaman Saini, Pradyumna K. Mishra, Asit R. Mridha, Rupesh K. Srivastava

**Author notes:** **<u>Send correspondence to:</u> Dr. Rupesh K. Srivastava,** Associate Professor, Department of Biotechnology, All India Institute of Medical Sciences (AIIMS), New-Delhi-110029, India.

## Abstract

Acute-respiratory-distress-syndrome (ARDS), the ultimate manifestation of acute-lung-injury (ALI) is a life-threatening respiratory failure with a significantly higher incidence and mortality worldwide. Recent discoveries have emphasized the existence of a potential nexus between gut and lung-health wherein physiology of the gut is directly linked with the outcomes of the lung pathologies. These discoveries fuel novel approaches including probiotics for the treatment of several respiratory disorders including ALI/ARDS. *Lactobacillus rhamnosus* (LR) is a preferred probiotic of choice as it has been reported to exhibit potent anti-inflammatory activities in various inflammatory diseases. In the present study, we investigated the prophylactic-potential of LR in lipopolysaccharide (LPS)-induced ALI/ARDS mice model, which mimics the pathophysiology of several respiratory disorders including respiratory tract infections, COVID-19, influenza, pneumonia, asthma, tuberculosis, cystic fibrosis, chronic obstructive pulmonary disease (COPD) etc. Our *in vivo* findings revealed that pretreatment with LR significantly attenuated lung inflammation and improved the pathophysiology of lung-tissues in ALI/ARDS mice. We observed that LR-administration suppressed the LPS-induced inflammatory cell infiltration in the lungs via ameliorating vascular-permeability (edema) of the lungs. Acute and chronic lung-disorders, including ARDS, are largely governed by innate-immune response. Interestingly, we observed that LR via modulating different ILCs (first responder to infections) subsets viz. ILC1, ILC2 and ILC3 prevent lung-fibrosis and maintain vascular permeability in LPS induced ALI/ARDS mice model. Of note, we observed a significant-enhancement in the percentage of inflammatory IL-17 producing CD3^-^Rorγt^+^NKp46^-^ ILC3-fraction along with a significant reduction in IL-22 (responsible for vascular integrity) producing CD3^-^Rorγt^+^NKp46^+^ ILC3 in both the BALF and lung-tissues. These ILCs would further augment the activation and recruitment of mononuclear phagocytic system (MPS-monocytes, macrophages and DCs) along with neutrophils and eosinophils in the lungs and BALF in ALI/ARDS mice model. Furthermore, gene expression and protein-analysis demonstrated that LR treatment significantly reduces the expression of inflammatory-cytokines in lung tissue and serum, thereby suggesting its potent immunomodulatory activity in attenuating ALI/ARDS. Summarily, our research convincingly establishes the prophylactic-role of LR in the prevention and management of respiratory-distress syndromes driven by the diverse inflammatory insults via modulating the lung’s “ILCs-MPS” axis with significant clinical-implications for the management of COVID-19, Influenza, COPD etc.

## Introduction

Acute respiratory distress syndrome (ARDS) was first described in humans nearly 50 years back and today, it is one of the most prevalent chronic disorders with escalating rates of disability and mortality worldwide. ARDS (severe form of acute lung injury) is one of the most difficult problems in critical-care medicine. It affects around a million patients annually and has a mortality rate close to 50% ^1^. The hallmarks in pathophysiology of ARDS are acute and widespread inflammatory damage to the alveolar-capillary barrier, increased vascular permeability, and accumulation of protein-rich edema fluid in the lungs which compromise gas exchange and result in hypoxemia ^2^. The major causes of ARDS can be grouped into following two categories: (1) direct pulmonary pathogenesis which includes bacterial and viral pneumonia, inhalation injury, aspiration pneumonitis, or trauma to the lung parenchyma, and (2) indirect extrapulmonary pathogenesis, such as extra-thoracic sepsis, trauma, shock, burn injury, transfusion, and other factors ^3^. Depending on the level of oxygenation, ARDS can be categorized into: ‘mild’, ‘moderate’ and ‘severe’. Lack of a definite diagnostic marker along with the disability to determine direct measurements of lung injury via pathological lung tissue samples in most patients make the diagnosis of ARDS solely based on clinical criteria. Furthermore, neither distal airspace nor blood samples can be used to diagnose ARDS.

An important factor involved in the pathogenesis of ARDS is the innate immune system. In ARDS, tissue injury is mediated by a network of immunological mechanisms involving various innate immune cells like ILCs, macrophages, dendritic cells (DCs) and neutrophils, Increased production of pro-inflammatory cytokines like interleukin (IL)-2, IL-6, IL-17, GM-CSF and IFN-γ further exacerbates the systemic inflammatory response and lung fibrosis which may be a pertinent factor in ARDS ^4^. The alveolar epithelium and vascular endothelium get affected as a result of inflammatory insults, either locally from the lungs or systemically from extra-pulmonary sites. These inflammatory insults further result in protein-rich edema fluid accumulation in the alveoli termed as “pulmonary edema” which compromises gaseous exchange and eventually leads to hypoxemia. Our immune system has developed to safeguard us against a variety of potential dangers without endangering the lung tissue. The primary function of monocytes, macrophages, and DCs collectively referred to as mononuclear phagocyte system (MPS), lining the respiratory tract, is to survey the lung microenvironment to distinguish between innocuous and harmful antigens and to elicit appropriate immune responses. Each immune cell type excels at a particular task: monocytes generate a lot of cytokines, macrophages have a high capacity for phagocytosis, whilst DCs are efficient at activating naive T cells. Numerous studies using murine models have established roles for the various populations of MPS both at steady state and during an infection or inflammatory response. Alveolar macrophages (AM) play a profound role in mediating inflammation as well as in the resolution of ARDS. Once AM are stimulated, they recruit circulating neutrophils and monocytes to the site of the lung injury and thereby initiate the inflammatory cascade ^5^. At steady state, lung contains a significant number of leukocytes, including specialized lymphoid cells known as innate lymphoid cells i.e., ILCs. ILCs are categorized into three subsets, ILC1, ILC2, and ILC3, which are thought to be the innate counterparts of Th1, Th2 and Th17 helper cells respectively. Although ILCs represent a small fraction of the pulmonary immune system, they are now considered as the first responders to the infection that further supports the establishment of a robust adaptive immune response. It is now also increasingly becoming clear that ILCs play a significant role in the emergence of various chronic lung illnesses ^6^. Recent studies revealed that alterations in the frequencies of ILCs are correlated with the disease severity in several respiratory diseases. However, the role of ILCs and its subsets in acute lung injury induced ARDS is still elusive. In the present study, we propose to investigate the role of “ILCs-MPS” network in ARDS pathophysiology.

Understanding the epidemiology, etiology, and pathophysiology of ARDS has come a long way in the last 50 years. Treatment majorly focuses on lung-protective ventilation since no specific pharmacotherapies have been identified till date. The most widely used adjuvant therapy in normal ARDS includes recruitment techniques, high-dose corticosteroids, and continuous neuromuscular blocking medications. Corticosteroids have anti-inflammatory properties and are observed to be a potential treatment modality for ARDS. Additionally, randomized trials to improve fluid treatment and mechanical ventilation for ARDS have shown promising clinical outcomes. Although several immunosuppressive drugs have been tested in clinical trials of patients with ARDS; however, almost none of them has proven beneficial in managing the disease in larger trials. The handful of them which showed promising outcomes is also associated with serious side effects. Thus, there is an exigent need for a potent immunomodulatory agent which can dampen the excessive inflammatory response in ARDS. Although supportive care for ARDS has greatly improved but unfortunately there are still no viable pharmacological treatments for ARDS. As a result, the search for safer and more affordable treatments having the capacity for both prevention and treatment of ARDS is still warranted. Experimental and clinical evidence demonstrates that gut microbiota possesses a crucial role in maintaining lung health. Recent research studies have suggested towards the existence of cross-talk between the gut and lung and this interplay has a major role in the control of infections ^7^. Many factors can alter the diversity and composition of the gut microbiota, leading to dysbiosis. Dysbiosis can also occur as a result of infections and chronic inflammatory or metabolic disorders ^8,9^. Changes in intestinal bacterial communities can influence disease outcomes even in distant organs (including the lungs), as demonstrated by transfer experiments with dysbiotic microbiota. The gut microbiota is dominated by the Firmicutes (e.g., Lactobacillus, Bacillus, and Clostridium), the Bacteroidetes (e.g., Bacteroides), and to a lesser extent, the Proteobacteria (e.g., Escherichia) and the Actinobacteria (e.g., Bifidobacterium) ^10^. Microbiota analyses in various respiratory infections have found a reduced abundance of the Lactobacillus genus (the Bacilli class) in the gut ^11^. Moreover, analysis of gut microbiome in studies involving COVID-19 patients has recently found that COVID-19 infection appears to blunt the growth of health-promoting gut bacteria such as Lactobacilli ^12^. Various studies have reported the protective role of probiotics including *Lactobacillus rhamnosus* (LR) in multiple respiratory and inflammatory diseases.

According to the Food and Agriculture Organization of the United Nations (FAO) and World Health Organization (WHO), probiotics are “living microbes that impart a health gain on a host when delivered in adequate amounts” ^13^. Our group has recently reported that LR via modulating the adaptive immune cells alleviates the inflammatory bone loss in osteoporosis ^13^. These findings suggest that probiotics have potential efficacy in dampening the severe inflammatory milieu in ARDS by modulating the immune system and thus would ameliorate respiratory immunity via modulating lung pathophysiology. Recently, a study reported that pre-treatment with *Lacticaseibacillus rhamnosus* significantly reduced the pathophysiology of ALI induced ARDS^14^. However, the potential of LR in dampening inflammatory milieu in ALI induced ARDS via modulating the “ILCs-MPS” network has not been deciphered yet. To our knowledge, this study for the first time will reveals the immunomodulatory potential of LR in significantly dampening the acute exudative phase (pulmonary edema and vascular permeability) of ALI/ARDS. The study thus proposes LR-administration as a plausible “prophylactic” therapy for managing ARDS in various lung pathologies including COVID-19, infectious disease etc.

## Materials and Methods

### Reagents and Antibodies

The following antibodies/kits were procured from eBiosciences (USA) and Biolegend (USA): BV605 Anti-Mouse-CD45-(103139), PerCp-Cy5.5 Anti-Mouse-CD11b-(45-0112-82), APC Anti-Mouse-Ly6G-(17-9668-80), BV421 Anti-Mouse-Ly-6C-(62-5932-82), BV786 Anti-Mouse-SiglecF-(78-1702-80), PE-Cy5 Anti-Mouse-CD11c-(15-0114-81), FITC Anti-Mouse-MHC Class II (I-A/I-E)-(107605), APC Cy7 Anti-Mouse-F4/80-(123118), staining buffer (0-5523-00) and RBC lysis buffer (00-4300-54). Lipopolysaccharides (LPS) Escherichia coli O111:B4 origin (L2630) and Evans blue (E2129) procured from Sigma, USA. *Lactobacillus rhamnosus* UBLR-58 was procured from Unique Biotech Ltd., Hyderabad, India.

### Animals

All in vivo experiments were carried out in 8-10 wks old male BALB/c mice. Mice were housed under specific pathogen-free (SPF) conditions at the animal facility of All India Institute of Medical Sciences (AIIMS), New Delhi, India and fed with sterilized food and autoclaved drinking water *ad-libitum*. Mice were randomly allocated into three groups with 6 mice in each group i.e., Control (mice received PBS); LPS (mice received LPS); and LPS + *Lactobacillus rhamnosus* (LR) (mice received LR for 14 days before LPS instillation). All the mice were kept in separate cages with light and dark periods of 12 hours each. All the procedures were performed as per the established animal ethical procedures of Institutional Animal Ethics Committee of AIIMS, New Delhi, India (382/IAEC-1/2022).

### Administration of Lactobacillus rhamnosus (LR)

Probiotic *Lactobacillus rhamnosus* (LR), UBLR-58 was purchased from Unique Biotech. Mice were orally gavaged with LR (10^9^ cfu) daily in 1X phosphate buffer saline for 2 weeks.

### Induction of ALI/ARDS with LPS

After two weeks of treatment, ARDS, and ARDS + LR group mice were anesthetized with ketamine (100 mg/kg) and xylazine (10-15 mg/kg) intraperitoneally and received an intranasal instillation of LPS (3 mg/kg). After 24 hours of LPS instillation mice were sacrificed and various organs harvested for further analysis.

### Collection of Bronchoalveolar Lavage Fluid (BALF)

For the bronchoalveolar lavage fluid (BALF) collection, tracheotomy was carried out and a cannula was inserted into the trachea after anesthetizing the mice with ketamine (100 mg/kg) and xylazine (10-15 mg/kg). Further lungs were lavaged three times with 1XPBS in a total volume of 3 ml (1 mL × 3). BALF was centrifuged at 1500 rpm for 10 min at 4 °C. The supernatant was aliquoted and stored in a freezer at −80 °C for cytokine analysis by ELISA and the cell pellet was suspended and fixed with 1X fixatives solution. The cell suspension was stored in dark at 4°C for further analysis by flow cytometry.

### Vascular Permeability assay

Pulmonary vascular permeability was assessed by Evans blue dye extravasation method. In brief, Evans blue dye (30 mg/kg) was given intravenously (i.v) to mice 60 minutes before the animals were euthanized. Mice were sacrificed, and both lungs were harvested. Left lung was cut into two halves and each half was weighed (wet weights). One half was dried in a drying oven at 150 ^0^C for 48 hours and the other tissue half was placed in 200µL formamide at 70^0^C for 48-72 hours. The concentration of Evans blue dye extracted in formamide was determined by spectrophotometry at a wavelength of 620 nm (BioTek Synergy H1). The dry tissue half which has been in the oven was weighed (dry weight). The dry/wet ratio of each lung sample was determined (index of edema) and used in the final calculation of Evans blue extravasation which was expressed as OD_620_/g of dry weight.

### Histological evaluation

For histological analysis, right lung was harvested, and a portion was cut and stored in 10% neutral formaldehyde for 24-48h. During the process, after following routine procedures for inclusion in paraffin, the lobes were next stained with haematoxylin-eosin (H&E). Analyses of histological sections were performed to determine the pulmonary changes induced by ARDS in the lung parenchyma. Lung injury was scored by a histopathologist as per the following criteria ^15^: pulmonary edema, vascular and alveolar features, and bronchiole pathology was graded: 0 (normal), 1 (mild), 2 (moderate), 3+ (severe). Points were added up and are expressed as median ± range of injury score.

### Flow cytometry

Cells were harvested from various tissues (BALF, lung and blood) and stained with antibodies specific for ILCs, macrophages, DCs, neutrophils and eosinophils. For the innate immune cells (Macrophages-AM & IM; DCs, Neutrophils, Eosinophils), cells were stained with anti-CD45-BV-605, anti-Ly6G-APC, anti-Ly6C-BV421, anti-SiglecF-BV786, anti-CD11b-PerCP-Cy5.5, anti-CD11c-PE-Cy5, anti-MHC Class II-FITC, and anti-F4/80-APC Cy7 antibodies for 45 min on ice in dark. For ILCs panel, cells were first stained with the anti-CD3-PE Cy7 antibody, incubated for 30-45 min on ice. After washing, fixation and permeabilization will be done, followed by incubation with fluorochrome labelled antibodies specific for ILC1, ILC2 and ILC3 were added viz. anti-Tbet-BV711, anti-GATA3-FITC, anti-Rorγt-PE and anti-NKp46-PerCp-Cy5.5 antibodies for 45 min on ice in dark. After washing, cells were acquired on BD-Symphony (USA). The samples were analyzed using Flowjo-10 (TreeStar, USA) software, and the gating strategy was carried out as per experimental requirements.

### Enzyme-linked immunosorbent assay (ELISA)

BALF was collected and centrifuged at 1500 rpm for 10 min at 4 °C, and then the supernatant was collected and stored in a freezer at −80 °C. Serum was collected from the blood and stored at at −80 °C till cytokine analysis. ELISA was carried out for the quantitative assessment of following cytokines i.e., IL-1β, IL-6, IL-17 and TNF-α in BALF and blood sera of all the mice groups by utilizing commercially available kits brought from BD (USA), as per the manufacturer’s instructions.

### RNA isolation

RNA was extracted from the lung tissue cells using the RNeasy Mini Kit (Qiagen, USA) according to the manufacturer’s instructions. The RNA quality was checked via 28s and 18s RNA using Bio Analyzer (Agilent Technologies, Singapore). Only samples with optimum RNA Integration Number value of ≥ 7 were used. RNA was also quantified using spectrophotometer (Nanodrop Technologies, USA). RNA was kept at −20 °C until further use.

### c-DNA Synthesis

For complimentary DNA conversion, 1µg of total RNA was used. First Strand c-DNA synthesis kit (Thermo Scientific, USA) was used according to manufactures instructions. The manufacturer’s recommended procedures for RT-PCR were followed, and c-DNA was kept at −20 °C until further use.

### q-PCR

Gene expression was measured using quantitative real-time (Applied Biosystems, Quantstudio^TM^-5, USA). Triplicate samples of cDNA from each group was amplified with customized primers viz. TNF-α (NM_013693.3), MCP-1 (NM_011333.3), IL-1β (NM_008361.4), IL-6 (NM_001314054.1), IP-10 (NM_021274.2), ROR-γt (AJ132394.1), IL-17A (NM_010552.3), IL-17F (NM_145856.2) and IL-22 (NM_016971.2) and normalized with arithmetic mean of GAPDH (NM_001289726.2) housekeeping gene. 25 ng of c-DNA was used per reaction in each well containing the 2X SYBR green PCR master mix (Promega, USA) along with appropriate primers. Threshold cycles values were normalized and expressed as relative gene expression.

### Statistical analysis

Statistical differences between the distinct groups were evaluated via student t-test paired or unpaired as required. All the values in the data are expressed as Mean ± SEM (n=6). Statistical significance was determined as p≤0.05 (*p ≤ 0.05, ** p ≤ 0.01, *** p ≤ 0.001, *** p ≤ 0.0001) with respect to the indicated groups.

## Results

### LR lowers pulmonary edema and maintains vascular permeability in ALI/ARDS

Firstly, we assessed the prophylactic potential of LR in mitigating ARDS in ALI mice model. For accomplishing the same, male BALB/c mice were randomly divided into three groups viz. control/healthy, ALI/ARDS and ALI/ARDS + LR. In ARDS + LR group, LR (10^9^ cfu) was administered orally for 14 days prior to LPS (3mg/kg) instillation. After 24hrs post-LPS instillation, mice were sacrificed and various tissues like blood, BALF and lungs were analyzed for different histological, vascular, and immunological parameters **(Fig. 1A).** We observed a significantly enhanced pulmonary edema (Lung wet/dry weight) in ALI/ARDS mice in comparison to the control group (p < 0.05), and LR administration significantly reversed the same in ALI/ARDS + LR group (p < 0.05) **(Fig. 1B-C).** To further examine the status of lungs vascular permeability in response to LR administration we performed Evans-Blue extravasation assay. Interestingly, we observed that LR was able to significantly reduce the pulmonary vascular leakage permeability (p ≤ 0.01) of ALI/ARDS + LR **(Fig. 1D).** These findings thus clearly indicate the potential of LR in attenuating acute exudative phase and improved the alveolar capillary permeability barrier in ALI mice model.

**Figure 1:**
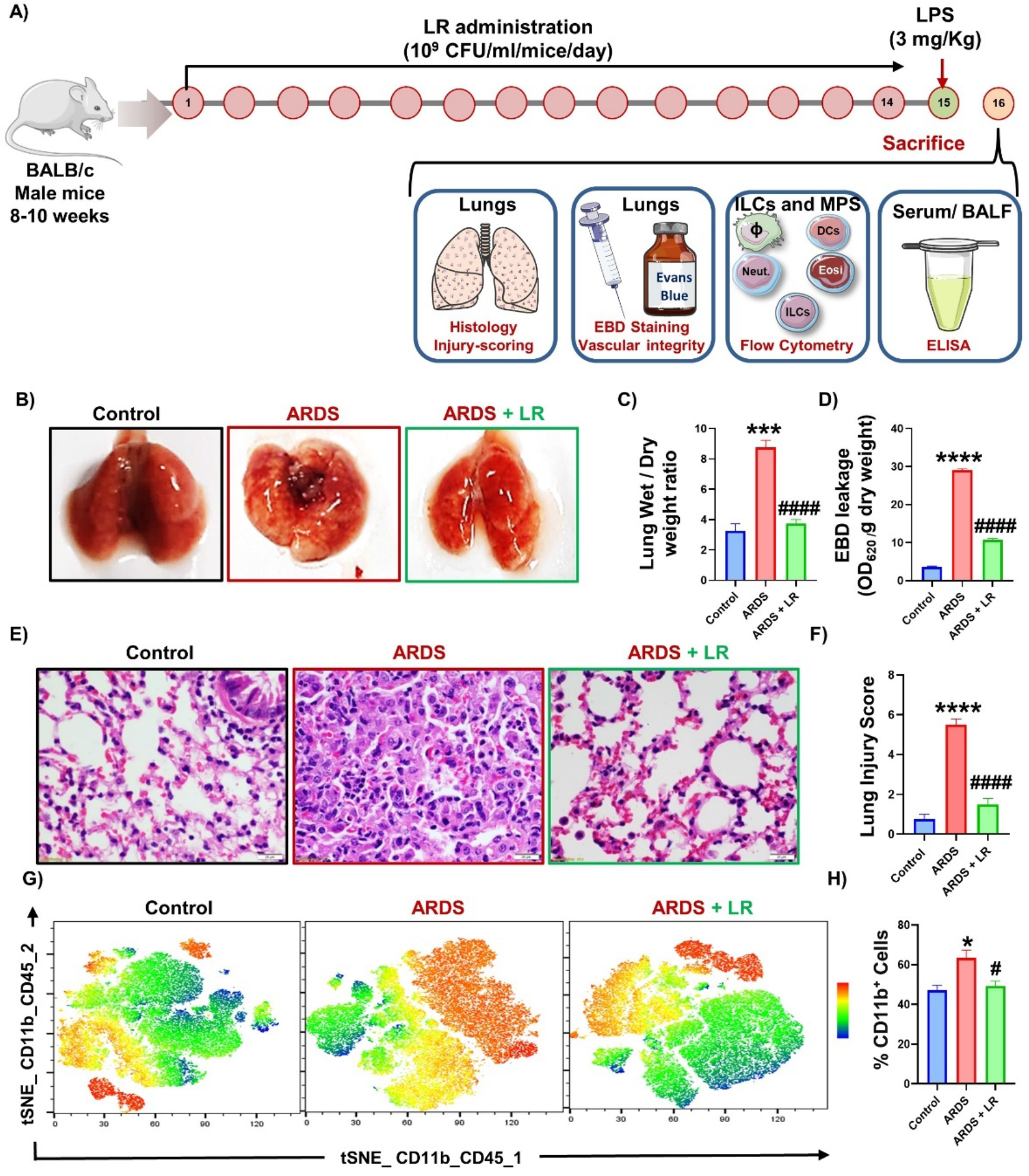
LR ameliorates pathophysiology of ALI/ARDS: Mice were divided into 3 groups, viz., control, ARDS, and ARDS+LR groups. ARDS + LR group received LR at 10^9^ CFU/day (100 μl) orally for 15 days. A) At the end of 15 days, LPS was given via intranasal route (3 mg/kg) and after 24 h mice were sacrificed, and various organs were harvested and analyzed for several immunological and histological parameters. B) Photomicrographs representing morphology of lungs. C) Bar graphs representing lung dry/wet weight ratio. D) Evans blue dye (EBD) leakage/dry weight. E) Photomicrographs were taken at 40X magnification after histological analysis by H & E staining. F) Bar graphs representing lung injury score. G) tSNE representing the cluster of CD11b^+^ cells in the CD45^+^ cell population. H) Bar graphs represents the percentage of CD11b^+^ cells in the lungs. Statistical significance was considered as (*p ≤ 0.05, **p ≤ 0.01, ***p ≤ 0.001) with respect to indicated groups (* indicated comparison between control and ARDS; # indicate comparison between ARDS and ARDS+LR).

Moving ahead, we next assessed the status of immune cells infiltration into the lungs in response to LR administration. Thus, we performed histological (H & E) staining and flow cytometry analysis to monitor the lung injury and cellular infiltration in ALI model. Our data showed significant increase in the immune cell’s infiltration, accumulation of fluid and alveolar hemorrhage in lungs parenchyma in ALI/ARDS group in comparison to the control group **(Fig. 1E).** In addition, lung injury score was observed to be significantly enhanced (5-fold; p < 0.001) in ALI/ARDS group with respect to the control group **(Fig. 1F)**, and LR administration significantly reversed these changes **(Fig. 1E-F).** Moreover, tSNE plot flow analysis revealed significant reduction in the CD11b^+^ cells (inflammatory marker) in LR group with respect to the ARDS group **(Fig. 1G-H)**. Altogether these results clearly suggest the immunomodulatory potential of LR in significantly improving the ALI/ARDS lung pathophysiology along with maintaining vascular permeability.

### LR administration ameliorates ALI-induced ARDS via modulating total ILCs

ILCs are important components of the pulmonary innate immune system as they are the first responders to any pathological challenge. Recent studies revealed that alterations in the frequencies of ILCs are correlated with the disease severity in asthma and COPD patients. But the role of ILCs in the pathophysiology of ALI/ARDS is still unreported. Thus, we were next interested in investigating the role of different subsets of ILCs in the pathophysiology of ALI/ARDS. For the same we isolated the blood, BALF and lung tissue cells from the lungs and analyzed them for various ILCs subsets via flow cytometry. Interestingly, our flow cytometric data revealed that the percentage of ILC1 (CD3^-^T-bet^+^) cells population was significantly enhanced in BALF (p < 0.01) and Lungs of ARDS group **(Fig. 2A-E)**, and LR administration significantly reversed these changes in BALF (p < 0.05) **(Fig. 2B-E).** Although the respiratory system contains all ILC subsets, but ILC2 play a critical role in lung homeostasis, repair/remodeling of damaged lung tissues, and the initiation of inflammation ^16–20^. Thus, we next investigated the status of ILC2 population in ALI/ARDS mice model and found that the frequencies of ILC2 (CD3^-^GATA-3^+^) cells population were significantly enhanced in the BALF (p < 0.001) and in the lungs (p < 0.05) **(Fig. 3A-E)** and administration of LR was able to significantly restore these trends in both BALF and Lungs **(Fig. 3B-E)**. Moreover, we observed that both the ILC1 and ILC2 cell population restored upon LR administration in the peripheral circulation **(Fig. 2F-G) (Fig. 3F-G).**

**Figure 2:**
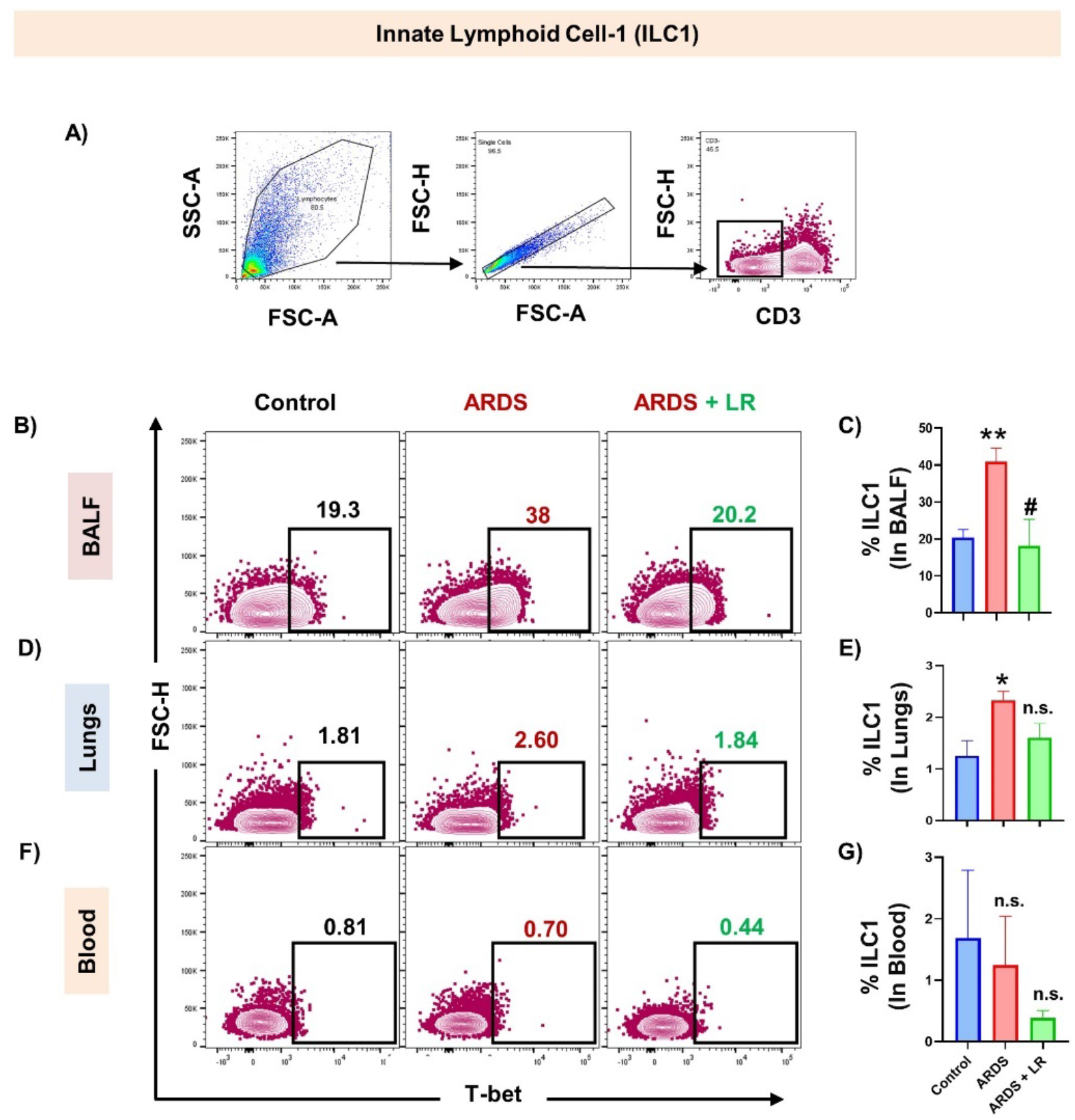
LR inhibits ILC1 (CD3^-^T-bet^+^) population: A) Gating strategy followed for CD3^-^T-bet^+^ ILC1 flow data analysis B) Contour plot representing percentage of ILC1 in BALF. C) Bar graphs representing percentage of ILC1 in BALF. D) Contour plot representing percentage of ILC1 in lungs. E) Bar graphs representing percentage of ILC1 in lungs. F) Contour plot representing percentage of ILC1 in blood. G) Bar graphs representing percentage of ILC1 in blood. Statistical significance was considered as (*p ≤ 0.05, **p ≤ 0.01, ***p ≤ 0.001, ****p ≤ 0.0001) with respect to indicated groups (*indicated comparison between control and ARDS; # indicate comparison between ARDS and ARDS+LR).

**Figure 3:**
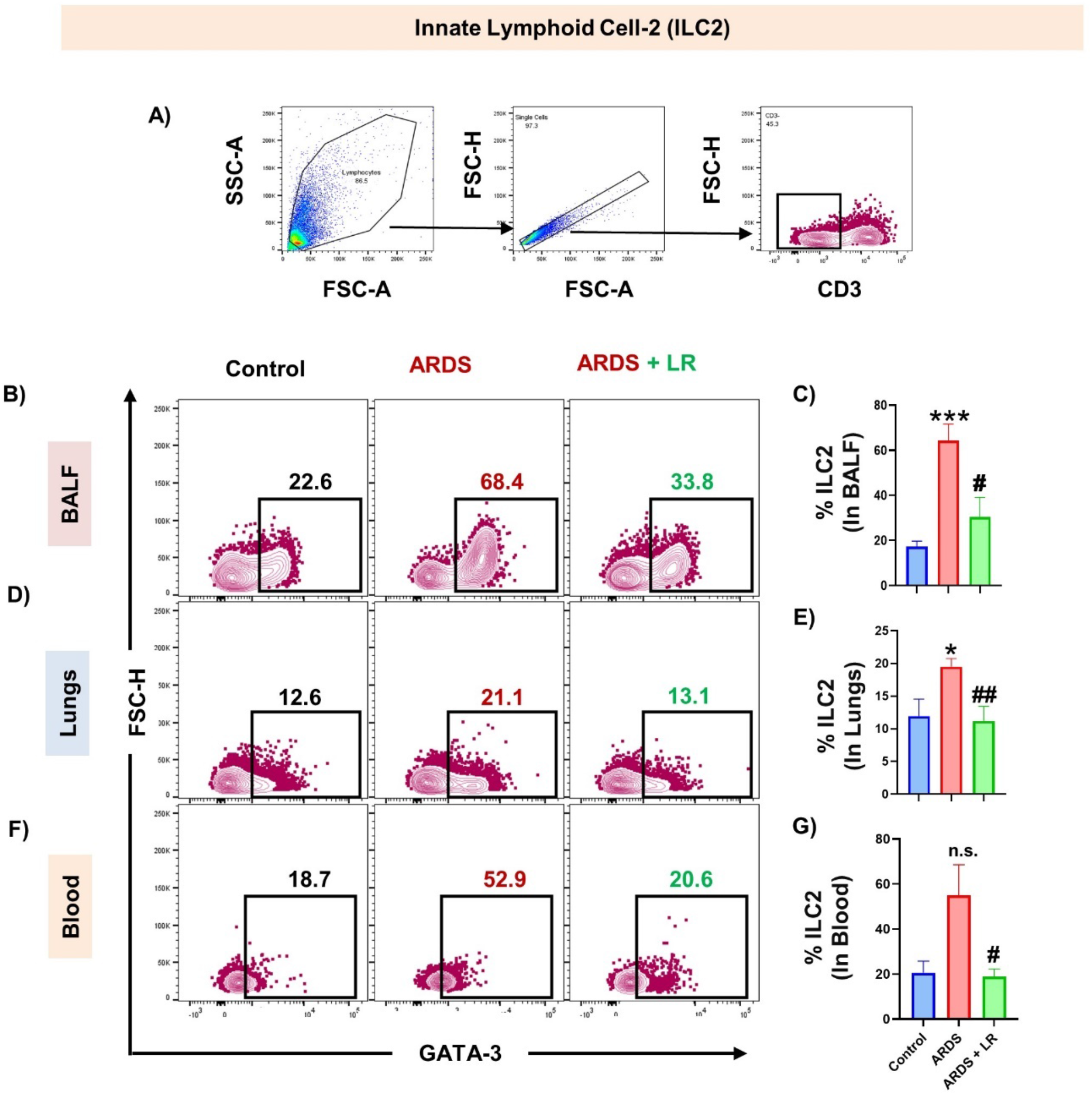
LR treatment modulates ILC2 (CD3^-^GATA-3^+^) population: A) Gating strategy followed for CD3^-^ GATA-3^+^ ILC2 flow data analysis B) Contour plot representing percentage of ILC2 in BALF. C) Bar graphs representing percentage of ILC2 in BALF. D) Contour plot representing percentage of ILC2 in lungs. E) Bar graphs representing percentage of ILC2 in lungs. F) Contour plot representing percentage of ILC2 in blood. G) Bar graphs representing percentage of ILC2 in blood. Statistical significance was considered as (*p ≤ 0.05, **p ≤ 0.01, ***p ≤ 0.001, ****p ≤ 0.0001) with respect to indicated groups (*indicated comparison between control and ARDS; # indicate comparison between ARDS and ARDS+LR).

The role of ILC3s in regulating mucosal homeostasis, inflammation and tissue remodeling has already been reported. Thus, we next investigated the role of ILC3 in the pathophysiology of ALI/ARDS. Interestingly, we observed that the frequencies of total ILC3 (CD3^-^Rorγt^+^) cells population was significantly enhanced in the BALF (p < 0.05) and administration of LR significantly reversed these changes but did not attained significance **(Fig. 4A-C).** However, we observed no modulation of total ILC3 cells in both the lungs and peripheral circulation of ARDS group in comparison to the control group **(Fig. 4D-G).** Our results thus clearly suggest the pivotal role of various ILCs subsets in the pathophysiology of ALI/ARDS.

**Figure 4:**
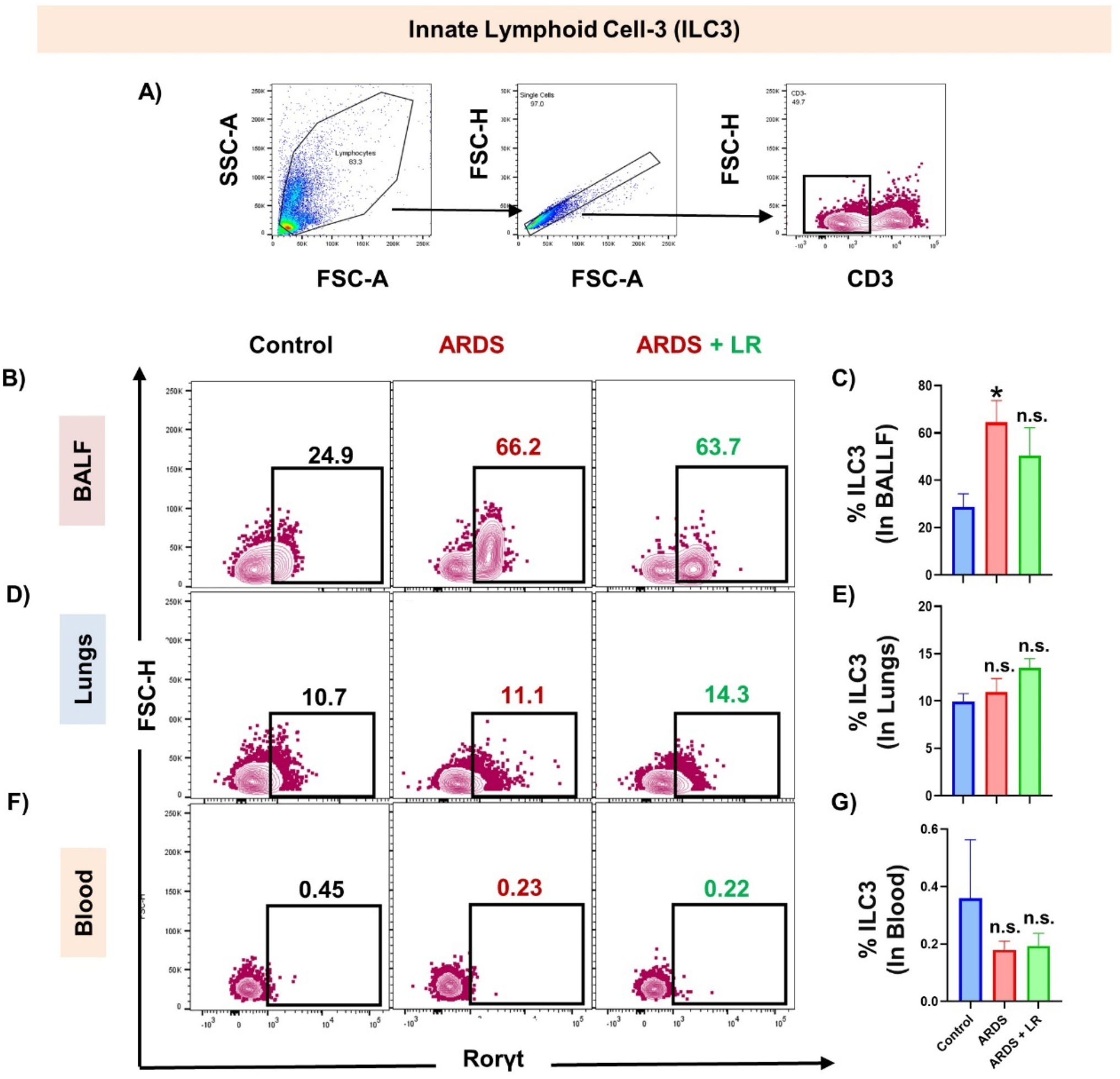
LR modulates ILC3 (CD3^-^Roryt^+^) population: A) Gating strategy followed for CD3^-^Roryt^+^ ILC3 flow data analysis. B) Contour plot representing percentage of ILC3 in BALF. C) Bar graphs representing percentage of ILC3 in BALF. D) Contour plot representing percentage of ILC3 in lungs. E) Bar graphs representing percentage of ILC3 in lungs. F) Contour plot representing percentage of ILC3 in blood. G) Bar graphs representing percentage of ILC3 in blood. Statistical significance was considered as (*p ≤ 0.05, **p ≤ 0.01, ***p ≤ 0.001, ****p ≤ 0.0001) with respect to indicated groups (*indicated comparison between control and ARDS; # indicate comparison between ARDS and ARDS+LR).

### NKp46^-^ILC3 and NKp46^+^ILC3 are pivotal in ARDS Pathophysiology

Based on the expression of NKp46, ILC3 subset can be further divided into two subpopulations viz. NKp46^-^ILC3 (IL-17 producing) and NKp46^+^ILC3 (IL-22 producing). To specifically examine which population contributes towards the pathophysiology of ALI/ARDS, flow cytometry was performed. Interestingly, we observed that in BALF, lung and blood the percentage of Rorγt^+^NKp46^-^ILC3 cells population was observed to be significantly enhanced in ARDS in comparison to the control group **(Fig. 5 A-C, E-F, H-I).** On the contrary, the percentage Rorγt^+^NKp46^+^ILC3 cells population was found to be significantly reduced in the BALF, blood and lung in ARDS group **(Fig. 5 D, G, J).** Interestingly, we observed that LR administration significantly reduced the percentage of Rorγt^+^NKp46^-^ILC3 cells population along with significantly enhancing the percentage of Rorγt^+^NKp46^+^ILC3 cells in the respective tissues (Fig. 5B-J). Moreover, we analyzed the expression of key genes associated with the ILC3 such *Rorγt, IL-17A, IL-17F* and *IL-22* among the different groups. Of interest, we observed that all these markers were significantly upregulated in ARDS as compared to the control and pre-treatment with LR significantly reduced the expression of these genes (Fig. 5 K-N). Altogether, our results for the first time delineate the roles of both NKp46^-^ILC3 and NKp46^+^ILC3 cell population in ALI/ARDS pathophysiology.

**Figure 5:**
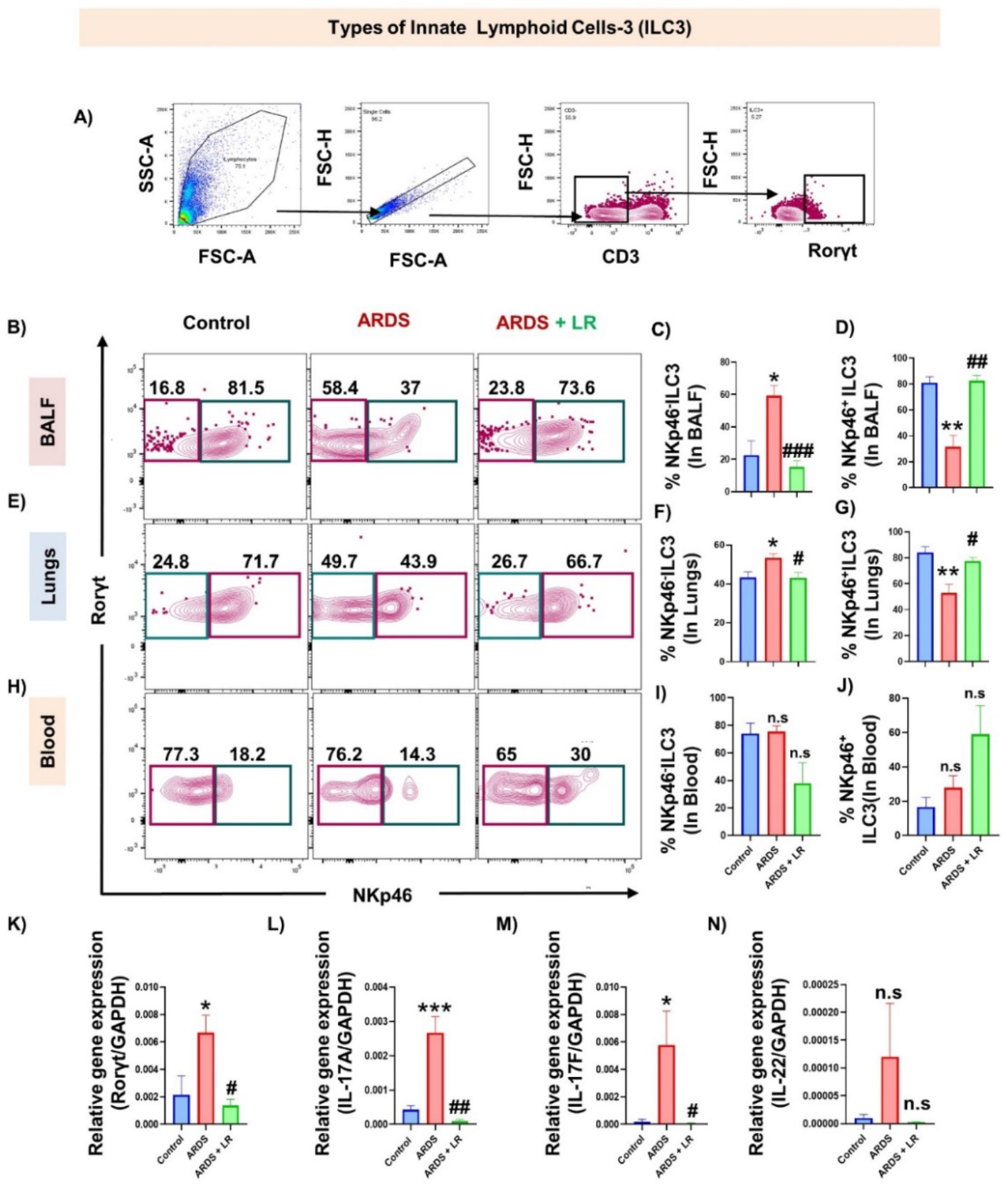
LR enhances CD3^-^Roryt^+^Nkp46^+^ILC3 and inhibits CD3^-^Roryt^+^Nkp46^-^ILC3 population: A) Gating strategy followed for CD3^-^Roryt^+^Nkp46^-^ILC3 and CD3^-^Roryt^+^Nkp46^+^ILC3 flow data analysis B) Contour plot representing percentage of CD3^-^Roryt^+^Nkp46^-^ILC3 and CD3^-^Roryt^+^Nkp46^+^ILC3 in BALF. C) Bar graphs representing percentage of CD3^-^Roryt^+^Nkp46^-^ILC3 in BALF. D) Bar graphs representing percentage of CD3^-^ Roryt^+^Nkp46^+^ILC3 in BALF. E) Contour plot representing percentage of CD3^-^Roryt^+^Nkp46^-^ILC3 and CD3^-^ Roryt^+^Nkp46^+^ILC3 in lungs. F) Bar graphs representing percentage of CD3^-^Roryt^+^Nkp46^-^ILC3 in lungs. G) Bar graphs representing percentage of CD3^-^Roryt^+^Nkp46^+^ILC3 in lungs. H) Contour plot representing percentage of CD3^-^ Roryt^+^Nkp46^-^ILC3 and CD3^-^Roryt^+^Nkp46^+^ILC3 in blood. I) Bar graphs representing percentage of CD3^-^ Roryt^+^Nkp46^-^ILC3 in blood. J) Bar graphs representing percentage of CD3^-^Roryt^+^Nkp46^+^ILC3 in blood. K) Relative expression of *Rorγt* gene. L) Relative expression of *IL-17A* gene. M) Relative expression of *IL-17F* gene. N) Relative expression of *IL-22* gene. Statistical significance was considered as (*p ≤ 0.05, **p ≤ 0.01, ***p ≤ 0.001, ****p ≤ 0.0001) with respect to indicated groups (*indicated comparison between control and ARDS; # indicate comparison between ARDS and ARDS+LR).

### LR modulates Lung Mononuclear phagocytic system (MPS) in ALI/ARDS

Mononuclear phagocytic system (MPS), comprising of macrophages, dendritic cells (DCs) and monocytes play a vital role in host defense and Lung tissue homeostasis ^21^. Thus, we next assessed the potential of LR in modulating the MPS system. Notably, we observed that the percentage of alveolar macrophages (AM) was significantly enhanced in ARDS group (p < 0.001) in comparison to the control group and administration of LR significantly reversed this trend (p < 0.01) **(Fig. 6A-C).** DCs serve a central role in inducing inflammatory immune response during infection; however, its fundamental mechanism in inducing pathogenesis of ARDS is still controversial. Notably, we observed that the percentage of DCs (CD45^+^LY6G^-^CD11b^+^SiglecF^-^LY6C^+^F4/80^-^CD11c^+^MHC-II^+^) were significantly increased in ARDS group whereas LR treatment significantly reduced the percentage of DCs in the lungs **(Fig. 6D-F),** thereby, suggesting towards the pathogenic role of DCs in ALI induced ARDS.

**Figure 6:**
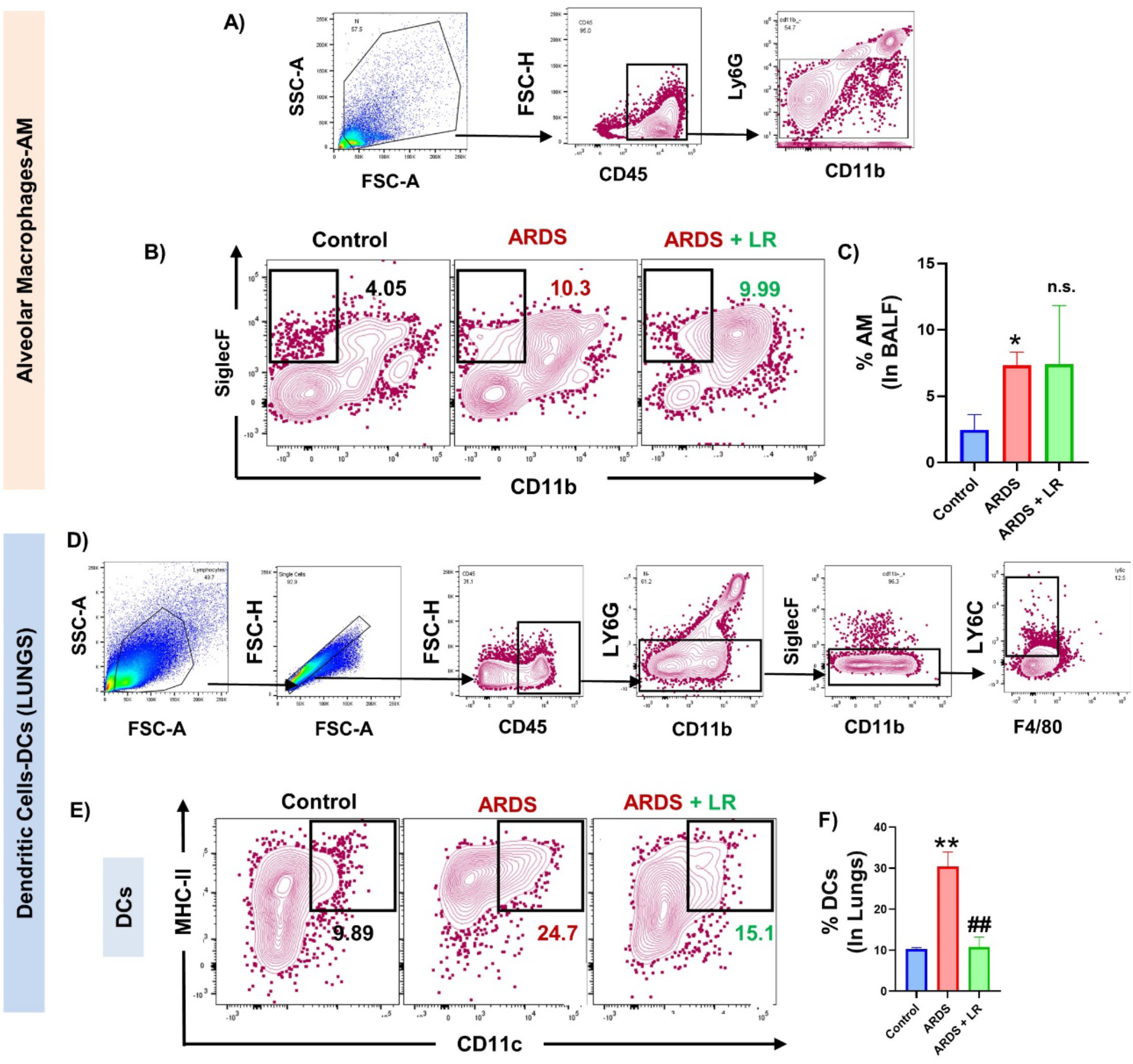
LR reduces Alveolar Macrophages (AM) (CD45^+^LY6G^+^CD11b^-^SiglecF^+^) and DCs (CD45^+^LY6G^-^ CD11b^+^SiglecF^-^LY6C^+^F4/80^-^CD11c^+^MHC-II^+^) in BALF and Lungs respectively: A) Representative image of gating strategy followed for BALF-AM (CD45^+^LY6G^+^CD11b^-^SiglecF^+^). B) Contour plot representing percentage of CD11b^-^SiglecF^+^ AM in BALF gated on CD45^+^LY6G^+^CD11b^+/-^ cells. C) Bar graphs representing percentage of percentage of AM. D) Representative image of gating strategy followed for lungs DCs (CD45^+^LY6G^-^CD11b^+^SiglecF^-^ LY6C^+^F4/80^-^CD11c^+^MHC-II^+^). E) Contour plots represent percentages of F4/80^-^CD11c^+^MHC-II^+^ DCs gated on CD45^+^LY6G^-^CD11b^+^SiglecF^-^LY6C^+^F4/80^-^ cells. F) Bar graphs representing percentages of DCs in lungs. Statistical significance was considered as (*p ≤ 0.05, **p ≤ 0.01, ***p ≤ 0.001, ****p ≤ 0.0001) with respect to indicated groups (*indicated comparison between control and ARDS; # indicate comparison between ARDS and ARDS+LR).

In addition, we further observed that the percentage of total interstitial macrophages (IM) were also significantly reduced in the ARDS group, however, no alteration was observed in the LR treated group **(Fig. 7A-C).** A study reported that IMs have three defined subsets viz. IM1, IM2 and IM3, which play an important role in the prevention of immune mediated allergic inflammation along with maintaining lung homeostasis. Importantly, we observed that the percentage of IM1 (CD45^+^LY6G^-^CD11b^+^SiglecF^-^Ly6C^int^F4/80^+^MHC-II^lo^CD11c^-^) were significantly reduced (p < 0.001), however the percentage of IM3 (CD45^+^LY6G^-^CD11b^+^SiglecF^-^Ly6C^int^F4/80^+^MHC-II^hi^CD11c^+^) were significantly enhanced (p < 0.01) in the lungs of ARDS group in comparison to control **(Fig. 7E, G).** Surprisingly, we observed that administration of LR significantly enhanced the percentage of IM1 (p < 0.01) along with simultaneously reducing the percentage of IM3 (p < 0.01) in ARDS group **(Fig. 7E, G).** Interestingly, we observed that the ratio of IM1 / IM3 was also significantly enhanced in the ARDS + LR group **(Fig. 7H).** However, no significant change was observed in the frequencies of IM2 population both in the ARDS and ARDS + LR group **(Fig. 7F).** In summary, our results for the first time clearly point to the prophylactic immunomodulatory potential of LR in ameliorating pathophysiology of ALI-ARDS via modulating the MPS in the lungs and BALF.

**Figure 7:**
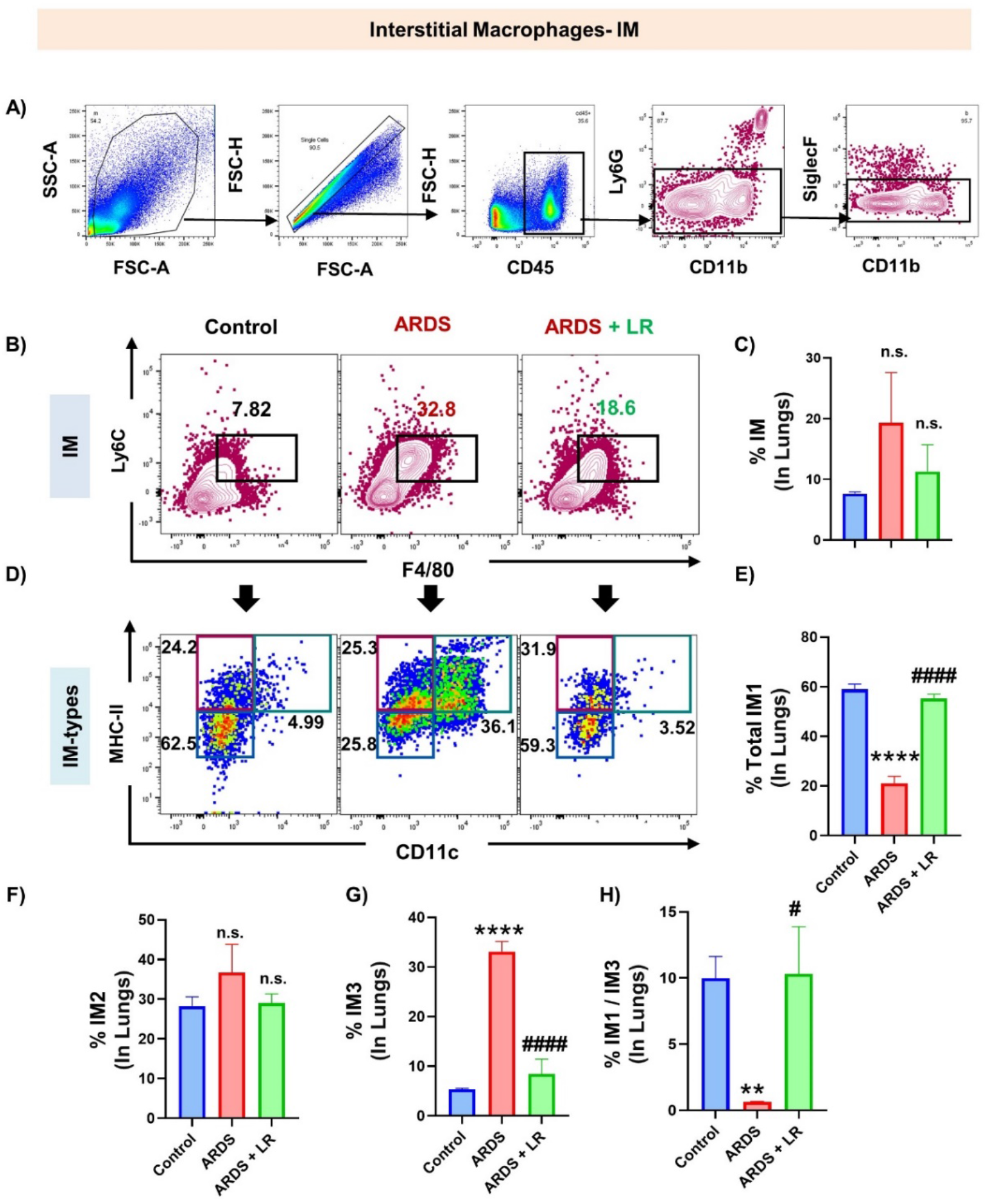
LR modulates interstitial macrophages (CD45^+^LY6G^-^CD11b^+^SiglecF^-^Ly6C^int^F4/80^+^) in lungs: A) Representative image of gating strategy followed for Interstitial macrophages-IM (CD45^+^LY6G^-^CD11b^+^SiglecF^-^ Ly6C^int^F4/80^+^). B) Contour plot representing percentage of IM in Lungs. C) Bar graphs representing percentage of IM. D) Plots representing IM1, IM2 and IM3 in lungs. E) Bar graphs representing percentage of IM1 in lungs. F) Bar graphs representing percentage of IM2. G) Bar graphs representing percentage of IM3. H) Bar graphs depicting IM1/IM3 ratio. Statistical significance was considered as (*p ≤ 0.05, **p ≤ 0.01, ***p ≤ 0.001, ****p ≤ 0.0001) with respect to indicated groups (*indicated comparison between control and ARDS; # indicate comparison between ARDS and ARDS+LR).

### LR modulates populations of Neutrophils and Eosinophils in ALI/ARDS

Acute lung damage is largely caused by activated neutrophils recruited to the lung tissues from the peripheral circulation. We thus next assessed the status of neutrophils in the BALF, lungs tissue cells and blood. Remarkably, our tSNE plot flow analysis showed that Ly6G expressing cluster of neutrophils was significantly enhanced in the ARDS group and LR administration was able to reduce the same **(Fig. 8A-B).** In consistent to this, the differential neutrophil count was observed to be significantly reduced in LR administered ARDS group in comparison to ARDS group (**Fig. 8C).** Also, our flow cytometric data further confirmed that the percentage of neutrophils (CD45^+^LY6G^+^CD11b^+^) were enhanced in BALF and lung (p < 0.01) (**Fig. 8D-G)** and LR administration significantly reversed these changes in BALF and lung (**Fig. 8D-G).**

**Figure 8:**
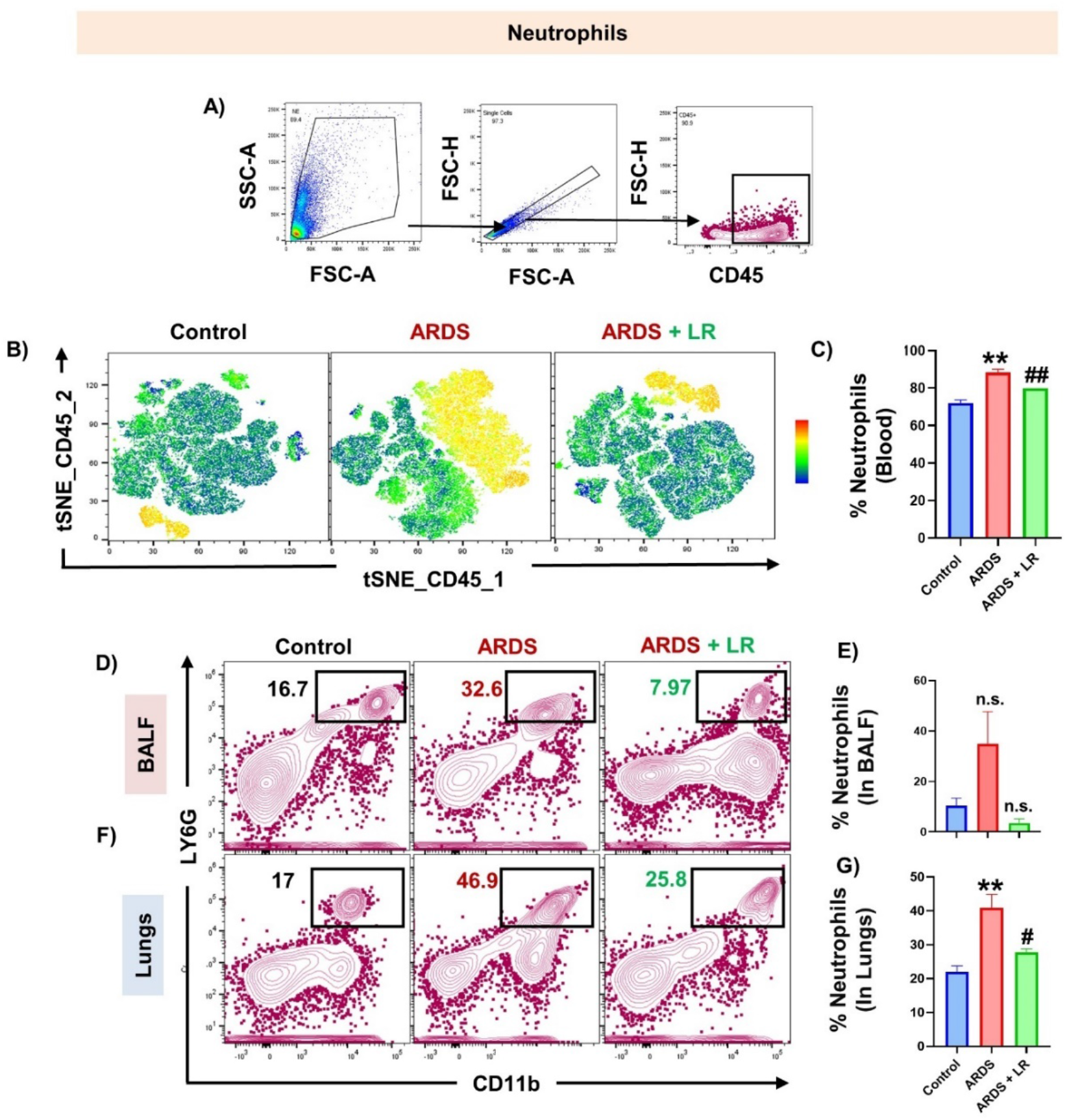
LR reduces Neutrophils (CD45^+^LY6G^+^CD11b^+^) population in BALF and lungs: A) Representative image of gating strategy followed for neutrophils (CD45^+^LY6G^+^CD11b^+^) in BALF and lungs. B) tSNE plot representing neutrophils population. C) Bar graphs representing percentages of neutrophils in blood. D) Contour plots representing percentages of LY6G^+^CD11b^+^ neutrophils gated on CD45^+^ cells in BALF. E) Bar graphs representing percentages of neutrophils in BALF. F) Contour plots representing percentages of LY6G^+^CD11b^+^ neutrophils gated on CD45^+^ cells in lungs. G) Bar graphs representing percentages of neutrophils in lungs. Statistical significance was considered as (*p ≤ 0.05, **p ≤ 0.01, ***p ≤ 0.001, ****p ≤ 0.0001) with respect to indicated groups (*indicated comparison between control and ARDS; # indicate comparison between ARDS and ARDS+LR).

Eosinophils play a significant role in type 2 inflammation but their role in the pathophysiology of ARDS is still unclear. Thus, we further looked into the status of the same in in the BALF, lungs tissue cells and blood. Strikingly, our tSNE plot flow analysis showed enhanced SiglecF expressing eosinophil cluster in the lungs of ARDS group **(Fig. 9A-B).** In consistent to this, we further observed enhanced eosinophils count, along with enhancement in the percentage of eosinophils (CD45^+^LY6G^-^CD11b^+^SiglecF^+^) in the BALF and lung of ARDS group **(Fig. 9C-G).** Remarkably, we observed that LR treatment reduced the eosinophils count in blood (**Fig. 9C)**. Altogether, these results establish the prophylactic potential of LR in the pathophysiology of ALI-ARDS via modulating the populations of neutrophils and eosinophils in the peripheral blood, lungs and BALF.

**Figure 9:**
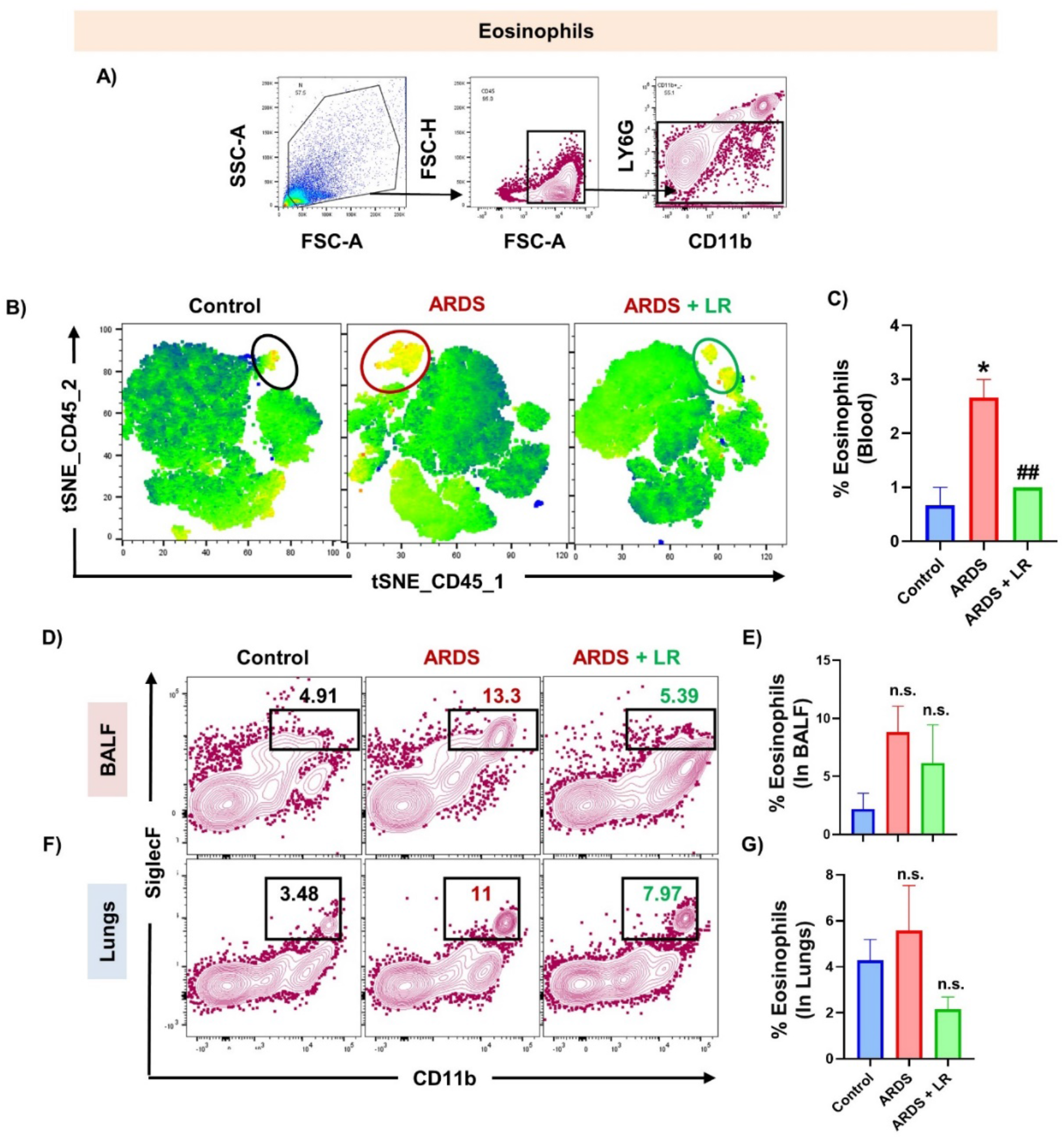
LR reduces Eosinophils (CD45^+^LY6G^-^CD11b^+^SiglecF^+^) population in BALF and lungs: A) Representative image of gating strategy followed for eosinophils (CD45^+^LY6G^-^CD11b^+^SiglecF^+^) in BALF and lungs. B) tSNE plot depicting the cluster of CD45^+^ population expressing SiglecF. C) Bar graphs representing percentages of eosinophils in blood. D) Contour plots representing percentage of LY6G^-^CD11b^+^SiglecF^+^ eosinophils gated on CD45^+^LY6G^-^ cells in BALF. E) Bar graphs representing percentages of eosinophils in BALF. F) Contour plots representing percentage of LY6G^-^CD11b^+^SiglecF^+^ eosinophils gated on CD45^+^LY6G^-^ cells in lungs. G) Bar graphs representing percentages of eosinophils in lungs. Statistical significance was considered as (*p ≤ 0.05, **p ≤ 0.01, ***p ≤ 0.001, ****p ≤ 0.0001) with respect to indicated groups (*indicated comparison between control and ARDS; # indicate comparison between ARDS and ARDS+LR).

### LR administration skews the expression of inflammatory cytokines and chemokines in ARDS

ALI/ARDS is a systemic inflammatory disease that affects the lungs along with various other systemic organs via inducing the expression of inflammatory cytokines. Markedly, we observed that the levels of inflammatory cytokines such as IL-1β (p < 0.01), IL-6 (p < 0.001), IL-17 (p < 0.01) and TNF-α (0.05) were significantly increased in both the serum and BALF of ALI/ARDS group in comparison to the control group **(Fig. 10A-B).** Of note, the expression of several inflammatory cytokine’s genes in lung tissue cells such as *IL-6, TNF-α, IL-1β* and *interferon gamma-induced protein-10 (IP-10)* also known as C-X-C motif chemokine ligand 10 (CXCL10) were observed to be significantly downregulated upon LR treatment in ARDS group **(Fig. 10C).** Moreover, the expression of monocyte chemoattractant Protein-1 (MCP-1/CCL2) was observed to be upregulated in the ARDS group, and LR treatment was able to significantly downregulate the expression of MCP-1 gene in ARDS group **(Fig. 10C)**, i.e., completely reversing the trend in ALI/ARDS + LR group, thereby suggesting the potent role of LR in attenuating cytokine-chemokine induced lung damage in the pathophysiology of ALI/ARDS.

**Figure 10:**
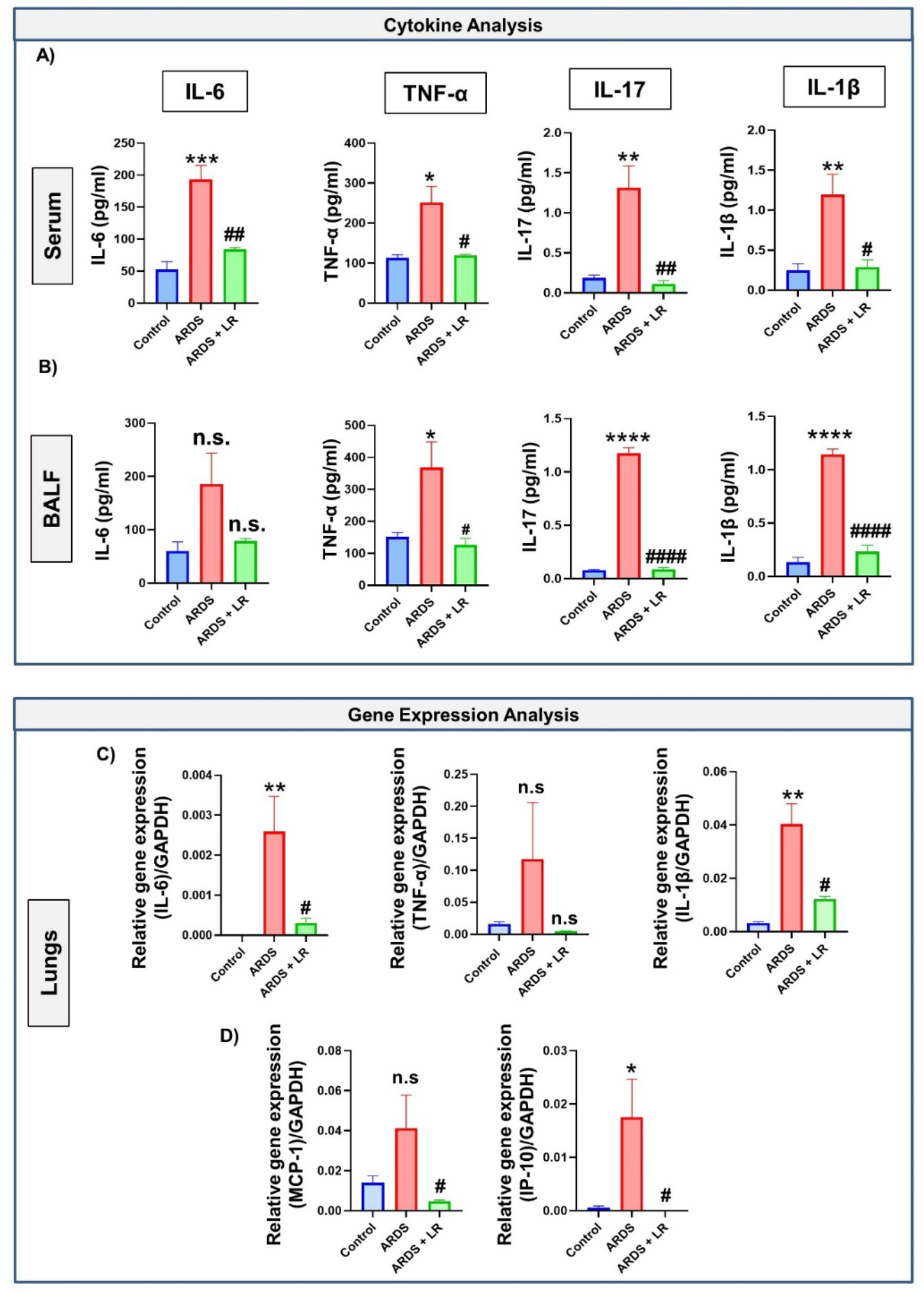
LR modulates the expression of cytokines in serum and BALF : A) Levels of inflammatory cytokines in sera. B) Levels of anti-inflammatory cytokines in BALF. C) Relative expression of *IL-6, TNF-α, IL-1β, MCP-1 and IP-10* genes. Statistical significance was considered as (*p ≤ 0.05, **p ≤ 0.01, ***p ≤ 0.001, ****p ≤ 0.0001) with respect to indicated groups (*indicated comparison between control and ARDS; # indicate comparison between ARDS and ARDS+LR).

## Discussion

The pathophysiology of ALI induced ARDS is initiated by various environmental challenges to the upper respiratory tract including viral (COVID-19, Influenza, Adenovirus, herpes simplex virus-HSV, cytomegalovirus-CMV), bacterial (*Mycobacterium tuberculosis*-M. tb), pollution (Allergens, Particulate Matter-PMs) etc. ARDS is a major cause of lung trauma, leading to severe morbidity and mortality. Diffuse interstitial and alveolar edema, inflammatory cell infiltration, and the production of proinflammatory factors are characteristic features of ALI, a severe lung inflammatory illness. An improved prognosis results from reduced alveolar inflammation and restored barrier function. The exudative phase of ALI/ARDS is characterized by recruitment of inflammatory cells, especially innate immune cells to the site of injury, which play a vital role for the host defense. ALI usually develops in patients with predisposing conditions that induce systemic inflammatory response among which sepsis is the major cause ^22^. Sepsis can occur primarily due to gram-negative bacteria in which LPS is the main component of their outer membranes ^23^. LPS is found to be the most important antigen that promotes the development of ALI ^24^. Therefore, the LPS-induced animal models provide novel ways in elucidating mechanisms for multiple diseases underlying ARDS. Moreover, Chen et al., reported that intranasal administration of LPS mimics the pathophysiology of human ALI/ARDS ^25^. Thus, in the present study, we employed LPS induced ALI/ARDS model to study the immunopathology of ARDS.

Probiotic strains including *Lactobacillus* sp., *Bifidobacterium* sp., and their metabolites have been reported to lessen the disease severity in tuberculosis, pneumonia and other respiratory viral infections ^26–28,26,29–31^. Recently, our group has too reported the immunomodulatory role of *Lactobacillus rhamnosus* in inflammatory disease ^13^. Experimental evidence suggests that administration of *Lactobacillus* can confer a beneficial role in various respiratory diseases including respiratory tract infections (RTIs), asthma, lung cancer, cystic fibrosis (CF) and COPD, thereby modulating respiratory immunity ^32–35^. Our group has also reported that LR administration skews the balance of inflammatory and anti-inflammatory cytokines and alleviates the inflammation induced bone loss in case of osteoporosis ^13^. In addition, a study reported that probiotics further lowers the oxidative stress rates and plasma peroxidation along with modulating cytokine balance ^36^. Strikingly, National Institute of Health (NIH) too is conducting more than 34 clinical trials to evaluate the efficacy of probiotics administration in COVID-19 patients (https://clinicaltrials.gov/). Together, these studies highlight the importance of probiotics in alleviating respiratory ailments. However, the potential immunological and cellular mechanism of LR in an ALI/ARDS model remains unclear. Thus, in the present study, we explored the immunomodulatory potential of LR as a prophylactic therapy in LPS-induced ALI/ARDS mice model.

Disruption of vascular integrity results in increased movement of proteins and fluid across the lung epithelium/endothelium. Moreover, there is accumulation of protein-rich inflammatory fluid in the normally fluid-free alveoli that results in increased lung weight. As a consequence of which, there is an increase in wet/dry lung weight ratio, an indicator of pulmonary edema. Our experimental data demonstrated that pulmonary edema is drastically increased in ARDS group with respect to control group. Interestingly, pre-treatment of LR significantly decreased the pulmonary edema as evidenced by reduced lung wet/dry lung weight ratio which is further indicative of reduced fluid accumulation in lungs and improved lung health. The protective effect of LR against lung vascular barrier disruption was further assessed by measurement of lungs vascular permeability. Our Evans blue exclusion data suggests that LR administration significantly reduced the ALI induced vascular permeability in lungs. Moreover, we further observed improved vascular integrity via significant reduction in the infiltration of immune cells along with reduced lung histological injury score in the lungs of ARDS + LR group with respect to ARDS group.

ILCs play an important role in tissue repair and homeostasis and are enriched at mucosal locations, such as the respiratory, gastrointestinal, and reproductive tracts, where they act as first responders to pathogens and assist both the innate and adaptive immune system to launch a quick defense. Recent studies demonstrated that enhanced frequencies of ILC1 in COPD patients correlated with the severity of disease and increased exacerbation risk and therefore suggest that ILC1 can be employed as a biomarker to monitor the progression of the disease. However, the role of ILC1 in acute exudative phase is still elusive. Our flow cytometric data revealed that the frequencies of ILC1 were significantly enhanced in BALF and lung tissues in ARDS group. Moreover, pre-treatment with LR significantly reduced the frequencies of ILC1 population in the respective tissues. In COPD patients it has been observed that alongside Th1 cells, ILC1 produce IFN-γ which via inducing elastolytic proteases and nitric oxide (NO) production by AM leads to pulmonary emphysema ^37^. Thus, we monitored the frequencies of AM in the inflamed lungs of ARDS group. We observed that in addition to the rise in ILC1, the frequencies of AM were also significantly enhanced in ARDS group in comparison to the control group. Interestingly, we observed that LR treatment was able to dampen AM population in ARDS group. This data was further supported by the downregulated expression of MCP-1 (regulate migration and infiltration of monocytes/macrophages) upon LR treatment. Thus, our data for the first time clearly highlights the pathogenic role of ILC1s and AMs in ALI/ARDS.

The pulmonary system contains all three ILC subsets, but the majority of research investigations so far has been focused on ILC2s, particularly in allergic airway inflammation (AAI) ^16,17,19,20,38^. In a model of cigarette smoke induced COPD, it has been observed that ILC2 population induce the recruitment of neutrophils, and its deprivation protected against the emphysema ^39^. Additionally, ILC2 are also known to promote migration of DCs and subsequent induction of Th2 cells ^17^. In asthma cases, it has been observed that ILC2 augments the recruitment of eosinophils via producing IL-4 cytokine. ILC2 via triggering aryl hydrocarbon receptor (AHR) can further promote mucus overproduction and disruption of the epithelial integrity ^40^. Together, these studies highlight the pathogenic role of ILC2s in various respiratory diseases. Nevertheless, its involvement in the pathophysiology of ARDS is still a matter of debate. In line with these studies, we thus observed a significant enhancement in the expansion of ILC2 population in our ALI/ARDS model and pretreatment with LR significantly reversed these changes.

ILC3s, are distinguished by the expression of the lineage-specific retinoic acid receptor-related orphan receptor-γt (RORγt). ILC3s are stimulated by myeloid or granulocyte cell-derived cytokines like IL-1β and IL-23, which cause them to release cytokines like IL-17A, IL-17F, IL-22, and granulocyte-macrophage colony-stimulating factor (GM-CSF). Moreover, rapid secretion of IL-17 and IL-22 highlights the crucial role of ILC3s in lung disease. Both these cytokines are necessary for the host’s defense against bacterial infection at mucosal surfaces, however, IL-17 primarily increases inflammation, whereas IL-22 typically acts as a tissue protector by preserving the integrity of the epithelial barrier, preventing lung fibrosis, and reducing airway inflammation^41,42^. In lung injury mice model, it has been observed that ILC3 is the predominant producer of proinflammatory IL-17A cytokine in the exudative phase (early) and contributes towards the pathogenesis via enhancing the recruitment of neutrophils. Further, IL-17 promotes the production of several inflammatory cytokines including TNF-α, IL-1β, IL-6, MCP-1 and GM-CSF ^43^. Interestingly, our flow cytometric data too indicated a significant enhancement in the percentage of ILC3 population in BALF of ARDS group and LR treatment significantly reduced the same. Excitingly, we further observed a significantly enhanced percentage of IL-17 producing NKp46^-^ ILC3 along with drastically reduced population of IL-22 producing NKp46^+^ ILC3 in the peripheral circulation, BALF and lungs of ARDS group. In addition, we observed that the levels of IL-17 cytokine were significantly enhanced in the sera and BALF of ARDS group. Also, we further observed downregulated expression of various inflammatory cytokine genes (*IL-17A* and *IL-17F*) and ILC3 associated transcription factor (*Rorγt*). Previous studies showed that ILCs population exhibits plasticity and ILC2 population can be converted into ILC1 and ILC3. Our data also demonstrates that ILC2 population are enhanced in BALF but reduced in lung tissue, thereby fueling the possibility that in ALI/ARDS, ILC2 are converted into either ILC1 or ILC3, but this needs further investigation. Altogether, our findings clearly suggest the pathogenic involvement of ILC1, ILC2 and ILC3 in the pathophysiology of ARDS, and pretreatment with LR was observed to significantly reduce this trend.

MPS detect danger signals through the pattern recognition receptors (PRR) and initiate innate responses. Activated AM and DCs further recruit effector cells for pathogen clearance and secrete first wave of cytokines including IFNα, IFNβ, IL-6, IL-12, and TNF-α to activate tissue resident lymphocytes ^44^. These cells further secrete second wave of cytokines that attract circulating neutrophils and monocytes, and stimulate AM to enhance phagocytosis ^44^. Based on these findings, we were thus next interested in evaluating the immunomodulatory potential of LR on the MPS network. Depending on the anatomical location, macrophages are considered as either AM and IM. Our flow cytometric data indicated that treatment with LR significantly decreased the population of inflammatory immune cells including neutrophils, AM, DCs and eosinophils. We observed that in ARDS, the percentage of IM were significantly reduced in ARDS group and LR treatment was unable to restore the reduced frequencies of IMs. Moving ahead, we further evaluated the distinct types of IMs viz. IM1, IM2 and IM3, and observed that the frequencies of IM1 were significantly reduced and IM3 were significantly enhanced in ARDS group. Our detailed analysis further revealed that pretreatment with LR significantly reversed both the frequencies and ratio of IM1/IM3 in ARDS + LR group. These data thus highlight the plausible pathogenic role of IM3 along with the disease reversing involvement of IM1 in the progression, development, and severity of ALI/ ARDS.

Moreover, we observed that via skewing the levels of inflammatory cytokines at both transcriptomic and protein levels such as IL-1β, IL-6, IL-17A, IL-17F and TNF-α, in both BALF and lungs tissues, LR was able to suppress the infiltration of innate immune cells and thus preserved vascular permeability of inflamed lungs in ALI/ ARDS. Altogether, these studies suggest that even a short-term (2 weeks) administration with probiotic-LR, significantly reduced pulmonary infiltration, vascular leakage, and cytokine release thereby ameliorating lung pathophysiology in ALI-ARDS via targeting the “ILCs-MPS” axis **(Fig. 11).** In conclusion, our results clearly demonstrate and establish that LR significantly attenuates the inflammatory responses and lung injury in LPS-induced ALI/ARDS mice. However, in the present study we only ventured onto the prophylactic role of LR and thus the therapeutic potential of LR still needs to be explored, thereby opening novel avenues for further research in the field.

**Figure 11:**
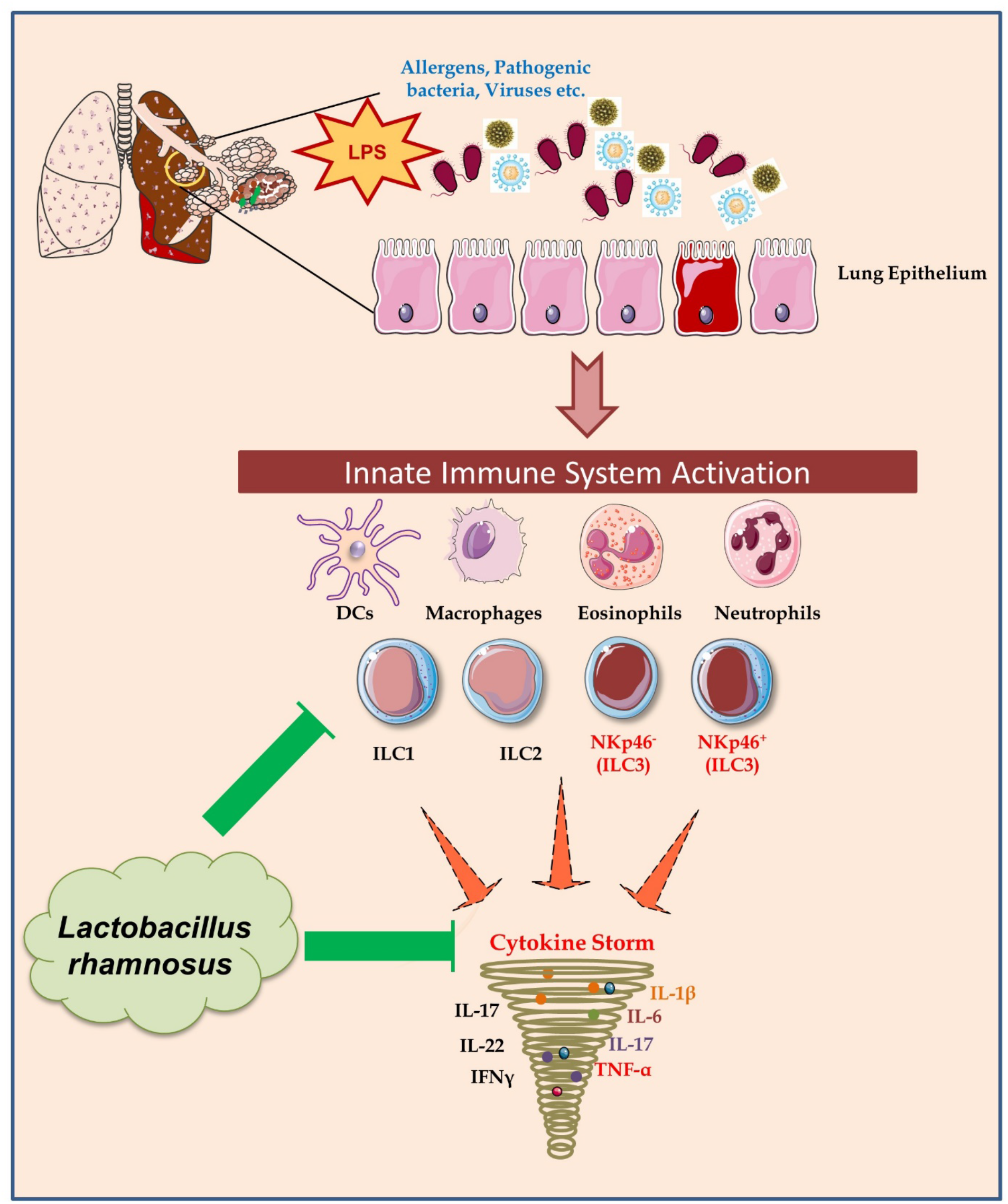
Summary of the results: LR via modulating the “ILCs-MPS” axis alleviates the LPS induced lung inflammation.

## Acknowledgement

This work was financially supported by projects: DBT(BT/PR41958/MED/97/524/2021), Govt. of India and Unique Biotech Ltd. (N-2312) sanctioned to RKS. LS, SD, CS and RKS acknowledge the Department of Biotechnology, AIIMS, New Delhi-India for providing infrastructural facilities. LS thank UGC for research fellowship. Figures are created with the help of Servier Medical Art, provided by Servier, licensed under a Creative Commons Attribution 3.0 unported license” (https://smart.servier.com).

## Author contributions

RKS contributed to the conceptualization and investigation of the study. LS, SD, CS, and RKS contributed to the methodology and formal analysis of data. PKM carried out cytokine analysis. RKS, LS, and SD contributed to the writing and editing of the manuscript. CS carried out gene expression analysis in the lung tissue cells. AR carried out histology and scoring of tissues. All the authors made a significant contribution in drafting, revising and critically reviewing the article for important intellectual content and gave final approval of the version to be published. All authors reviewed the manuscript.

## Compliance with ethical standards

All applicable institutional and/or national guidelines for the care and use of animals samples were followed.

